# Optimising Genotype Imputation for Precise Genetic Association in Forensic Phenotype Prediction and Trait Studies

**DOI:** 10.1101/2025.08.01.668059

**Authors:** Zehra Koksal, Andreas Tillmar

## Abstract

The imputation of single nucleotide polymorphisms (SNPs) provides a low-cost alternative to augment the size of genotyped SNP panels. Genotype imputation is commonly applied to study genotype-phenotype correlations in medical and population genetics, and has a great – yet unexplored – potential in a forensic context. Forensic DNA phenotyping, i.e., the prediction of phenotypic traits based on SNPs, can greatly benefit from imputing missing DNA markers necessary for utilising available prediction models and implementing novel prediction models. Currently however, most imputation studies investigate the performance of random SNPs with limited focus on SNPs involved in phenotypic expression or association.

In the current study, individuals from the 1000 Genomes Project with high predicted trait diversity were used to explore the imputation accuracy of SNPs leveraged in phenotype prediction models and SNPs associated with facial traits compared to all SNPs. Further, the performance of the HIrisPlex-S prediction model for phenotypic traits was investigated using different imputed datasets.

Firstly, we were able to corroborate that the number and selection of SNPs in the genotype dataset and the minor allele frequency (MAF) are major drivers of imputation call and error rates. Secondly, we explored increased imputation errors for phenotypic SNPs compared to randomly selected SNPs due to MAF differences. Further, we corroborated findings on lower imputation error rates for SNPs in coding regions due to increased linkage compared to non-coding regions. When investigating the impact of imputation on the performance of trait prediction using the HIrisPlex-S prediction model, we observed that datasets with more genotyped SNPs and phenotypes with more observations in the reference panel improved the prediction of these phenotypes. Finally, we showed novel insights into the improved trait prediction when applying more lenient calling thresholds for SNP imputation due to the detrimental impact of missing genotypes on trait prediction accuracy compared to imputation errors.

Our findings, which show different imputation performances for general compared to phenotype-associated and prediction-model SNPs, highlight the importance of investigating imputation performances for the SNPs of interest. Further, we reported optimal trait predictions using lenient calling threshold of imputed SNP genotypes paired with a SNP panel with high linkage, which shows the high applicability of SNP imputation for phenotypic trait predictions. We recommend imputation tests for the prediction models of interest due to the differences between prediction models.

**Highlights:**

- Number and selection of SNPs in imputation input and MAF impact call and error rate
- Lower imputation accuracy of phenotype-associated SNPs compared to random SNPs
- Lower imputation error rates for SNP in coding over non-coding regions
- High abundance of phenotype in reference panel favours its prediction
- Most accurate trait predictions for lenient SNP calling thresholds for imputation

## 1. Introduction

A low-cost alternative to augment the size of an available single nucleotide polymorphism (SNP) panel is the imputation of SNPs. This in-silico method predicts alleles of loci that are absent in the dataset obtained from in-vitro experiments (i.e., in test samples), but that are present in a reference panel. Imputation tools utilize allele information on phased sequences (i.e., haplotypes) of the test sample to find the most probable shared segments (that are identical by descent) between the test and reference panel samples. Within shared segments, ungenotyped alleles in the test sample can be filled from the shared segment of the reference haplotype. This is possible because SNPs on shared segments are non-randomly associated (linkage disequilibrium LD) and thus the SNP allele information on such a haplotype implies the presence of additional SNP allele information on the same segment (1,2).

The benefit of genotype imputation lies in its potential to study variants that have not directly been genotyped, but can also be used to correct and verify genotype calls (6), which is called “genotype refinement”. The applicability of genotype imputations depends on its success imputing the true missing genotypes (3–5). Thus, previous studies have identified several factors impacting the imputation performance and recommendations to optimize the imputation.

The size and composition of the reference panel has a great impact on the imputation accuracy. Larger panels tend to improve imputation accuracy due to increased probability to observe SNP patterns matching the test samples (1,2). Most importantly, the imputation will benefit from a reference panel comprised of similar individuals as the test samples. This has recently prompted the implementation of customized population-specific or population-inclusive reference panels (7–10). Secondly, increased number of SNPs in the test samples has shown to increase imputation accuracy, as more sites can match between test and reference panel samples (1,2). Naturally, inaccuracies in the test and reference panel samples affect the imputation accuracy (2). A study comparing the effect of drop-in and drop-out error types on the imputation accuracies, showed that both error types resulted in fewer imputed alleles and higher imputation error rates when compared to no introduced errors. Particularly, drop-in error types (turning homozygote genotypes to heterozygote genotypes) introduced the biggest imputation error rates (11). Finally, rare alleles with low minor allele frequencies (MAFs) are observed less often in the reference panel, reducing the chance to find other SNPs linked with the rare allele, thus making them harder to impute (1,2).

Currently, genotype imputation is widely utilized to study a larger number of genetic variants with an increased power, e.g., by combining datasets from different SNP panels in meta-analyses (3–5). Particularly rare variants may profit from this approach. Another major benefit is the increased chance to detect causal SNPs associated with a trait, making imputation a crucial part of genome wide association studies (GWAS) (12–16). However, when validating imputation performances, generally all SNPs are considered (1,7,8,11,13,14,23). Alternatively, for applications addressing certain traits, validating imputation performances specifically of phenotype-associated SNPs (12,15–19) could be more representative, as these SNPs are mostly restricted to certain genomic regions. The genomic regions affect the size of LD blocks, which has a central role in the imputation performance (1,2,24,25). Typically, LD blocks are more prominent in regions that escape recombination (24,26), (recently) underwent natural selection (26–29), translate to proteins or functional molecules (genic regions) (28), and carry lower frequently-common SNPs (MAF:10-20%) (25,28).

Overall, SNP imputation finds increasing applications in population and medical genetics of humans (7,8,17–19) and non-human species (20–22), but its potential in forensic genetics has only recently begun to be explored (11,30). Genotype imputation can compensate for DNA markers missing due to the crime scene samples’ low DNA quality and quantity or targeted genotyping (i.e., from forensically validated multiplex genotyping and targeted massively parallel sequencing approaches) (30,31). In cases where routine identification analyses cannot be applied, it is helpful to provide an investigative lead to narrow down the potential sample donors (31). These can be obtained from one or several alternative forensic genetics strategies, such as familial searches or forensic DNA phenotyping (FDP). FDP utilizes genetic markers to predict phenotypic traits (i.e., externally visible characteristics (EVCs), biogeographic ancestry (BGA) and age) (31). The applicability of genotype imputation for FDP, and the factors influencing its effective use, remain uninvestigated, although genotype imputation promises to be very impactful for FDP. Since especially rare variants benefit from imputation due to an increased statistical power (3–5), imputation increases the chance of correctly predicting less frequent phenotypic traits, which are more informative investigative leads compared to common traits. Further, genotype imputation has the potential to catalyse trait prediction by increasing the statistical power in GWAS, thereby increasing the number and effect sizes of SNPs associated with phenotypic variance, which is essential for accurate prediction models (31).

In the current study, we explore the imputation of SNPs connected with various phenotypic traits of the human face. For this, we compare the imputation performance for SNPs identified in GWAS studies (phenotype-associated SNPs “PhenoSNPs“) and SNPs selected for phenotype prediction models (phenotype prediction model SNPs “PredSNPs“) in comparison with all SNPs (“AllSNPs“). The comparison is based on 31 selected individuals from the 1000 Genomes Project of diverse biogeographic ancestries and diverse combinations of phenotypic traits according to our predictions. Finally, we assess the impact of imputation parameters on the accuracy of phenotype prediction models.

## 2. Material and Methods

### 2.1 Data collection and selection

To test the imputation performance particularly for SNPs involved in phenotype expression, test samples of diverse phenotypic traits were selected. For this, (a) hair, eye and skin colours were predicted using the HIrisPlex-S prediction model (32–34) and (b) biogeographic ancestry clusters were estimated using the software STRUCTURE (35).

Prior to applying HIrisPlex-S and STRUCTURE, phased variant data of 2,504 unrelated individuals from the 1000 Genomes Project phase 3 dataset (36) were downloaded. For prediction (a), 39 out of the 41 SNPs used in the HIrisPlex-S prediction model (missing SNPs: rs312262906_A and rs201326893_A) were extracted and the phenotype probabilities were estimated using the HIrisPlex-S webservice (https://hirisplex.erasmusmc.nl/, accessed 05.03.25). The most probable traits per category were estimated using decision trees in accompanying papers (32,37).

For prediction of biogeographic ancestry clusters (b), STRUCTURE v2.3.4 (35) was run for all 2,504 samples using a burn-in of 10,000 and number of MCMC repetitions of 20,000 for 4 to 6 clusters (K) in 4 independent runs each. The most fitting number of clusters K was determined applying the software STRUCTURE harvester v0.6.93 (38) on all STRUCTURE output files. CLUMPP v1.1.2 (39) was applied using the largeKgreedy algorithm to combine the populations and individuals STRUCTURE output files for the most probable K.

For the following analyses, the most diverse set of individuals was selected by picking 1 individual per combination of predicted traits from (a) the HIrisPlex-S model. For phenotypes seen in more than 1 individual, the selection criterion extended to individuals of (b) diverse biogeographic ancestry coefficient clusters. This resulted in a set of 31 phenotypically diverse individuals.

### 2.2 Removal from reference panel

For imputation and data quality tests, the 31 individuals were removed from the phased reference panel of 2,504 1000 Genomes Project samples, resulting in a reference panel of the remaining 2,473 individuals.

### 2.3 Data cleaning

The 1000 Genomes Project variant data of the 31 individuals were processed to remove sites with multiple entries of the same genomic locus with differing reference and alternative allele entries hinting towards ambiguous sites. Structural variants affecting more than 2 nucleotides (as reference or alternative allele) were removed, thus only retaining single nucleotide variants and short insertion and deletions. For conservative variant filtering, the program conform-gt (version from 24.05.2016, https://faculty.washington.edu/browning/conform-gt.html#introduction) was applied to the vcf files of the 31 individuals alongside the reference panel of 2,473 individuals to ensure that the vcf entries are consistent with the reference panel. Variants flagged as “ambiguous” or “failing” by conform-gt were removed from the vcf files, if the reference allele and/or any of the alternative alleles did not match 1 of the 4 possible nucleotides.

### 2.4 SNP genotype imputation and performance statistics

For imputation performance tests, 7 subsets of the multisample vcf file of the 31 individuals were generated, representing the observed genotype data (from here on referred to as “preimputation datasets“). The preimputation datasets were filtered to contain either 70%, 30%, 10%, 5% or 1% randomly selected loci of the multisample vcf file for each chromosome 1 to 22 and chromosome X. Further, 2 preimputation datasets were generated to include only the 677,864 SNPs of the Infinium Global Screening Array-24 v3.0 BeadChip (40) (from here on referred to as “AncestryDNA panel“) and the 5,446 SNPs of the FORCE panel (41). These panels were selected due to their relevance and frequent application in genetic research (42), biobank studies (43), pharmacogenomics (44), consumer genomics companies (40), and in forensic and investigative settings (45–47).

These 7 preimputation datasets were used as input data for data imputation using BEAGLE v5.5 (1) with the following parameters: burnin=3, iterations=12, phase-states=280, imp-states=1600, imp-segment=6, imp-step=0.1, imp-n steps=7, cluster=0.005, ap=true, gp=true, ne=1,000,000, window=40, overlap=4.0 and the HapMap GRCh37 as used map. A custom python script was written to call genotypes according to a defined genotype or allele probability threshold of 0.95 and 0.99. Failure to meet the genotype or allele probability threshold resulted in non-called genotypes.

To quantify the imputation performance, call rates describe the fraction of imputed and observed SNP genotypes divided by all SNP genotypes in the reference panel. The genotype error rate was calculated as the fraction of incorrectly imputed SNP genotype count divided by the sum of preimputed (observed) and imputed SNP genotype count.

### 2.5 SNP identification and characterisation

The imputation performance of SNP genotypes connected with phenotypic traits were explored. For this, 1,164 unique SNPs were identified that were utilized in 23 phenotypic trait prediction models (32,48–64) (“PredSNPs“; Supplementary Table 1). Secondly, 4,663 unique SNPs were identified that were associated with phenotypic traits reported in the GWAS catalogue (accessed 20.02.2025) applying a list of 29 keywords (Supplementary Table 2 and 3) and in single GWAS publications (15,48) (“PhenoSNPs“).

The imputation performance of SNP genotypes in coding versus noncoding regions was tested after establishing if single genomic loci corresponded to the loci in the GENCODE database (version 37 for lift GRCh37).

### 2.6 Predicting HIrisPlex-S phenotypes based on imputed datasets

The HIrisPlex-S predictor was used to predict the most probable eye, hair and skin colour per sample as described above based on the imputation datasets using genotype probability thresholds 0.99, 0.95, 0.9, 0.85, 0.5 and 0.2.

## 3. Results and Discussion

In the present study, we tested the imputation performance of SNPs connected with phenotypic traits. SNPs that are present in the 1000 Genomes Project dataset and leveraged in phenotype prediction models are referred to as “PredSNPs” and SNPs associated with phenotypic traits as “PhenoSNPs”. For comparison, all SNPs in the 1000 Genomes Project dataset are referred to as “AllSNPs”.

For these 3 SNP sets, we tested the impact of different imputation parameters and the number of SNPs in the preimputation dataset (i.e., observed genotype data) on the number of final SNPs and their error rates. The tested imputation parameters included genotype and allele probability thresholds with cut-offs 0.95 and 0.99. The SNP density over all chromosomes in the preimputation dataset ranged from 30% missing data (21,424,640 SNPs) to 99% missing data (306,056 SNPs) including the 2 SNP panels AncestryDNA of ∼98% missing data (618,383 SNPs) and the FORCE panel of ∼99.9% missing data (4,612 SNPs).

### 3.1 Number and selection of SNPs in preimputation dataset and minor allele frequency of SNPs are major drivers of call and error rate

As shown in Figure 1, a general trend shows that fewer SNPs in the preimputation dataset resulted in reduced call rate. Since fewer SNPs from the test samples can be identified in the reference panel, the number and size of linkage blocks with co-inherited SNPs is reduced as well. These are expected observations seen in the literature (11). Exceptions to this trend are presented by the AncestryDNA preimputation dataset, which covers a broad range of carefully selected genome-wide and clinically associated SNPs. Importantly, the panel includes human linkage markers with high informative value for a wide range of linked markers as tested in imputation studies of the commercial vendor (40). Hence, the informative value per SNP is higher in the AncestryDNA panel than in the preimputation datasets with randomly selected SNPs. Thus, the AncestryDNA panel reaches a higher call rate despite the lower number of SNPs prior to imputation.

**Figure 1:**
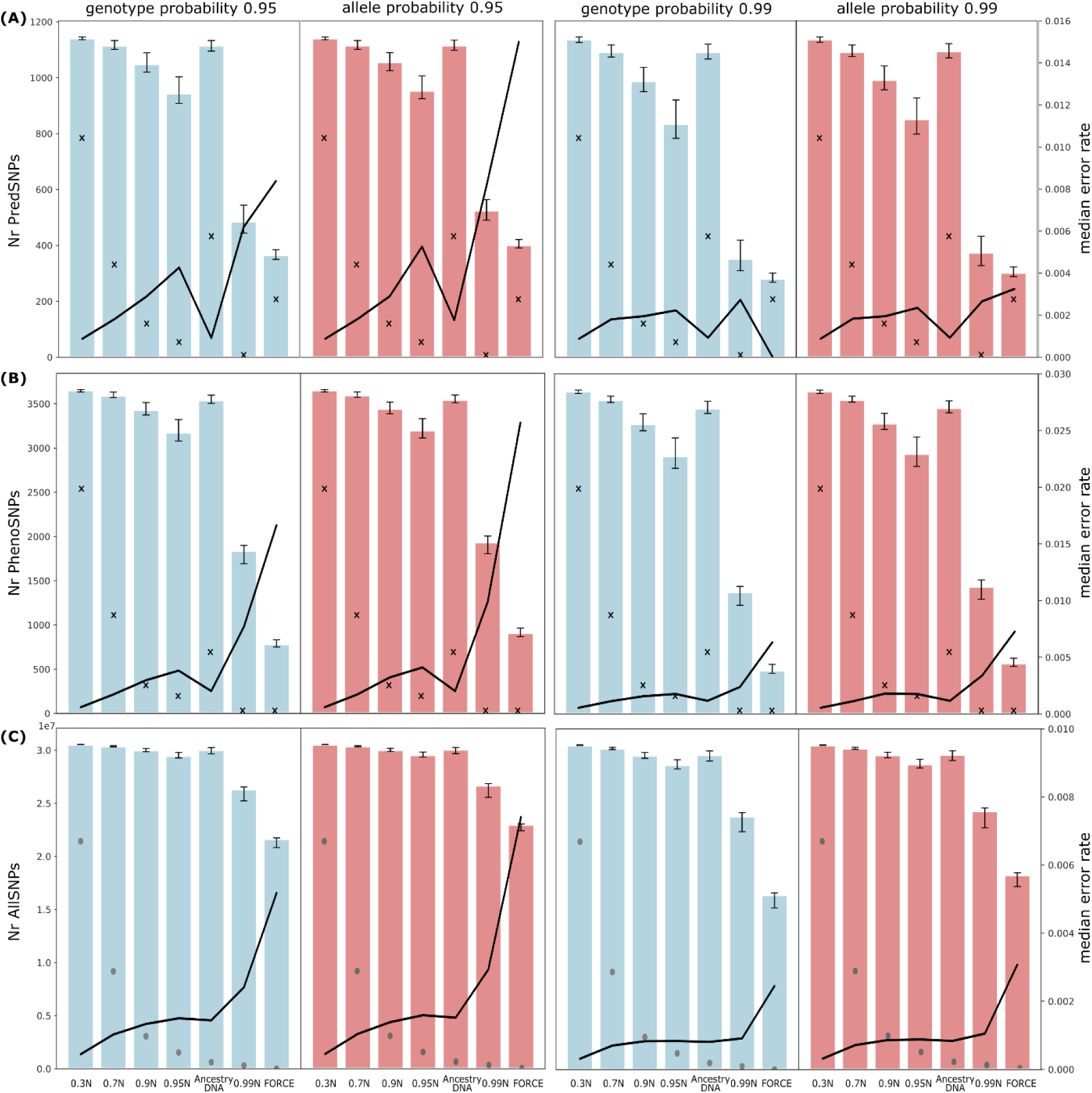
Imputation performance for (A) SNP genotypes from phenotype prediction models (PredSNPs), (B) SNP genotypes associated with phenotypic traits (PhenoSNPs), (C) and all imputed SNP genotypes (AllSNPs). SNP genotypes were imputed in sets of the same 31 samples with increasing fractions of missing data from left to right on the x axes. The preimputation datasets correspond to missing data fractions of 30% (21,424,640 SNPs), 70% (9,181,980 SNPs), 90% (3,060,653 SNPs), 95% (1,530,320 SNPs), ∼98% (AncestryDNA panel: 618,383 SNPs), 99% (306,056 SNPs) and ∼99.9% (FORCE panel: 4,612 SNPs). The number of SNPs in the preimputation datasets are marked as grey dots in subplot (C). Alleles were called using probability thresholds of 0.95 (left) or 0.99 (right) for genotypes (blue) or alleles (red). Line graphs corresponding to the secondary axes show median genotype error rates over all called SNPs. Black crosses in subplots (A) and (B) correspond to the number of PredSNPs and PhenoSNPs present prior to imputation.

Generally, the error rate increases with decreasing number of SNPs in the preimputation datasets (Figure 1), because lower SNP densities correspond to fewer “anchoring” SNPs in fewer and smaller linkage blocks in the reference panel. This causes correctly imputed SNP genotypes to drop more than the total imputed SNP genotypes, which increases the error rate.

The negative correlation between SNP density in the preimputation dataset and error rate is also observed in findings from the literature (11), where the error rates are slightly higher than that of AllSNPs in the current study (Figure 1C; Supplementary Figure 1). These slight differences can be caused by different levels of similarity between sample and reference panel and by differently filtered SNPs (see Data Cleaning under Material and Methods).

The AncestryDNA panel presents an exception to the SNP density and error rate trend, due to the previously described high informativeness of its SNPs. Further, imputations with a strict genotype probability threshold of 0.99 for the PredSNPs (Figure 1A) using the FORCE panel resulted in an exceptionally low error rate (median: 0). This observation is confounded by the FORCE panel composition, which already includes 203 PredSNPs. For a genotype probability threshold of 0.99, only 77 SNPs were imputed when presenting the median value per sample. This immensely reduces the risks of observing wrongly imputed SNP genotypes.

Further, when the number of SNPs in the preimputation dataset decreases, the median MAF of the imputed SNP genotypes decreases as well (Figure 2, solid line). This is because SNPs with lower MAFs have major alleles with more observations in the reference panel, compared to SNPs with higher MAFs with more even observations of the major and minor alleles. This increases the imputation chances of low-MAF SNP genotypes. Accordingly, among the imputed SNP genotypes, those that are incorrectly imputed tend to have higher MAFs (Figure 2, dashed line).

**Figure 2:**
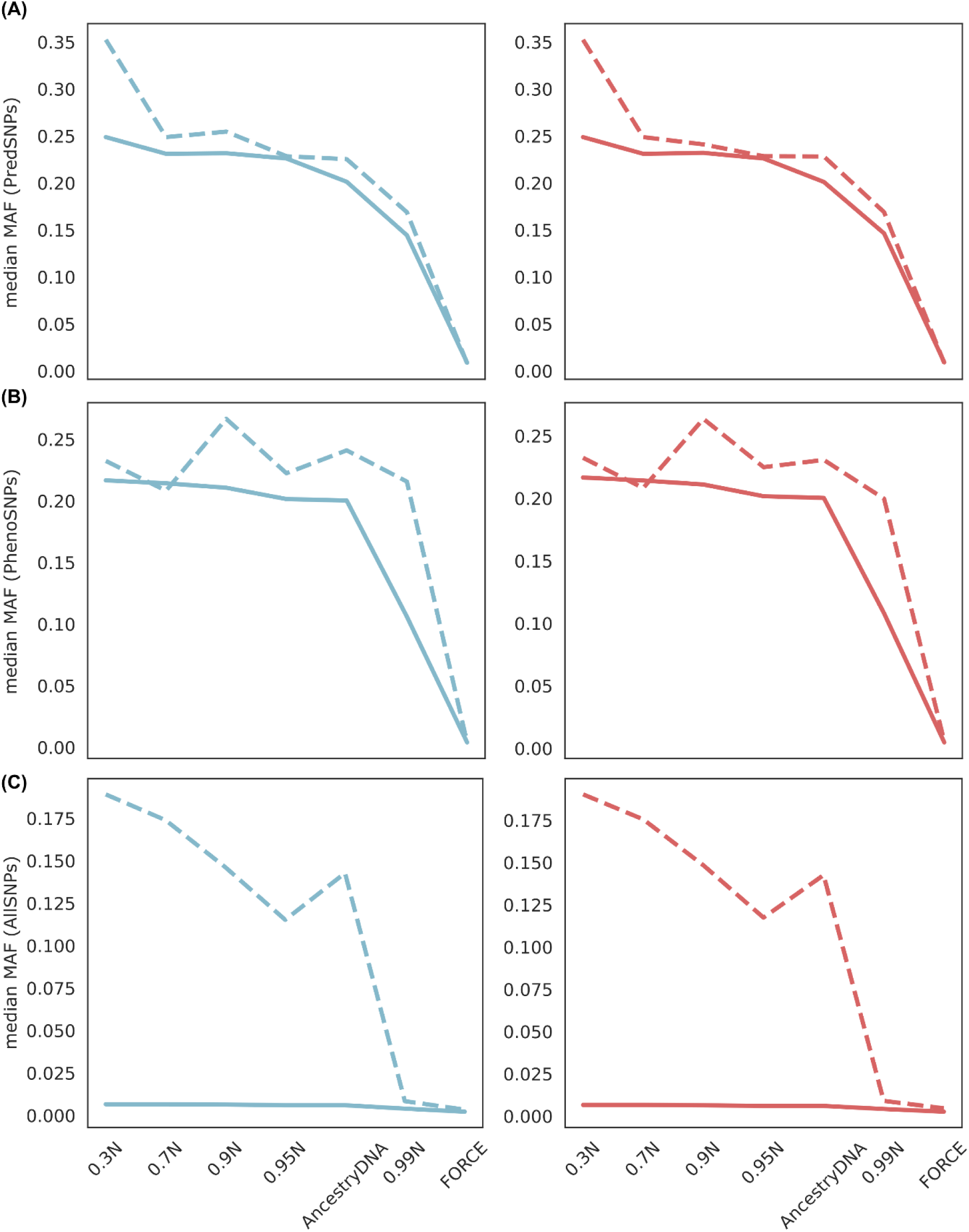
Median minor allele frequencies (MAFs) of wrongly and correctly imputed SNP genotypes (continuous line) and only wrongly imputed SNP genotypes (dashed line) for (A) PredSNPs, (B) PhenoSNPs, (C) AllSNPs, when using genotype probability (blue) or allele probability (red) cut-off 0.99.

Finally, the positive correlation between MAF and error rate is supported by the lower error rates for AllSNPs (median MAF: 0.00699) compared to PredSNPs and PhenoSNPs (median MAFs: 0.240815 and 0.215855, respectively) (Supplementary Figure 2).

### 3.2 PredSNPs and PhenoSNPs tend to be more often incorrectly imputed compared to randomly selected SNPs due to MAF differences

Given the 10,000 to 26,000 times lower number of SNPs of the PredSNPs and PhenoSNPs compared to AllSNPs, we further explored imputation error rates after normalising the number of randomly selected SNPs in imputed datasets. We randomly selected SNPs in equal number to those exceeding the genotype probability cut-off of

0.99 for PredSNPs and PhenoSNPs, respectively. As presented in Figure 3, the median error rates of the same number of randomly selected SNPs are generally slightly lower for random SNPs than for the PredSNPs and PhenoSNPs. These differences are statistically significant as shown using a Wilcoxon signed-rank test to compare randomly selected SNPs with PredSNPs (p=0.03125 when removing FORCE panel; p=0.296875 if retaining FORCE panel) and PhenoSNPs (p=0.015625 for all panels). As described above, this most likely is the consequence of the MAF differences, which were on median 0.006789 (range=0.00639-0.007188) for the randomly selected SNPs, while PredSNPs and PhenoSNPs had median MAFs of 0.226038 (range=0.009185-0.248802) and 0.201677 (range=0.004792-0.216653) for the 7 different preimputation datasets (Figure 3).

**Figure 3:**
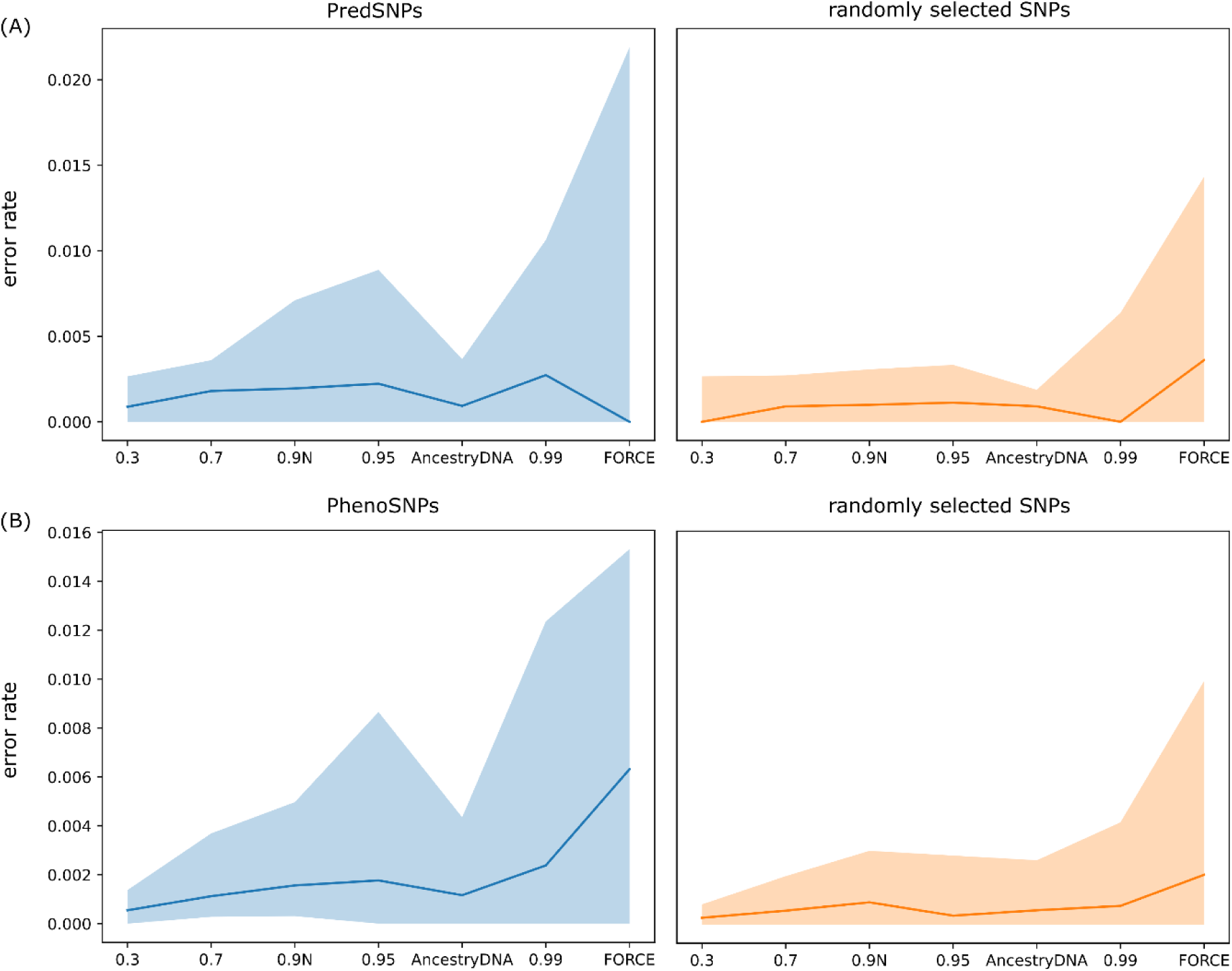
Comparison of error rates of PredSNPs (A) and PhenoSNPs (B) (blue) to an equal number of randomly selected SNPs (orange) for 7 preimputation datasets (x axes). Median error rates are presented as dark blue or orange lines with the areas from the lowest to highest error rates represented in lighter shades.

As seen before in Figure 1, Figure 3 shows that the more informative SNPs in the AncestryDNA panel resulted in reduced error rates for phenotype-correlated (PredSNPs and PhenoSNPs) and randomly selected SNPs. However, the error rate reduction in the AncestryDNA panel relative to the other preimputation datasets is more drastic for phenotype-correlated SNPs (Figure 3, left) compared to randomly selected SNPs (Figure 3, right). This is likely due to the composition of the AncestryDNA panel, which includes more of the phenotype-correlated SNPs (see crosses in Figure 1 A, B).

While the differences in MAFs explain the observed data, the question arises whether the increased error rates for PredSNPs and PhenoSNPs are also impacted by differing error rates between SNPs of coding and non-coding regions.

### 3.3 Lower imputation error rates for SNP genotypes in coding regions compared to non-coding regions due to increased linkage in coding regions

To test for differences in imputation accuracy between SNP genotypes on coding and non-coding regions, we identified SNPs of both regions on a random chromosome (chromosome 16) representative of the whole genome, using the GENCODE database (version 37 for lift GRCh37). The MAFs of SNPs on the coding (median=0.006589, average=0.062084) and non-coding (median=0.006589, average=0.062823) regions were near-identical and their impact on observed trends negligible.

The number of SNPs on coding regions were ∼3.3 times higher (chr16=754,304) than those on non-coding regions (chr16=228,560). This trend remained consistent for the 7 different preimputation datasets. To account for the uneven number of coding and non-coding region SNPs when calculating error rates, the number of SNPs on coding regions after imputation were subsampled to the number of non-coding SNPs for each sample and preimputation dataset.

As seen in Figure 4, the error rates for SNPs on non-coding regions were generally higher than for SNPs in coding regions. For the 3 largest preimputation datasets with 30%, 70% and 90% missing data, this trend was statistically significant (Mann-Whitney U test p-values 3.4077e-03, 2.6596e-02 and 7.1665e-03). This aligns with findings in the literature, where lower imputation error rates were reported for SNPs in coding regions compared to SNPs in intronic and particularly intergenic regions (65). One major driver of this trend is the increased linkage between SNPs in coding regions compared to non-coding regions (28,65). Because selection leads to more constraints for functional alleles, the allele and nearby hitchhiking SNPs result in more co-inherited SNPs and thus larger blocks in LD (28). This leads to the increased recombination observed outside of genes, as shown for individual chromosomes (66). For the smallest preimputation dataset, the FORCE panel, this trend was not observed, possibly due to too few SNPs in the preimputation dataset to result in a meaningful difference between coding and non-coding regions.

**Figure 4:**
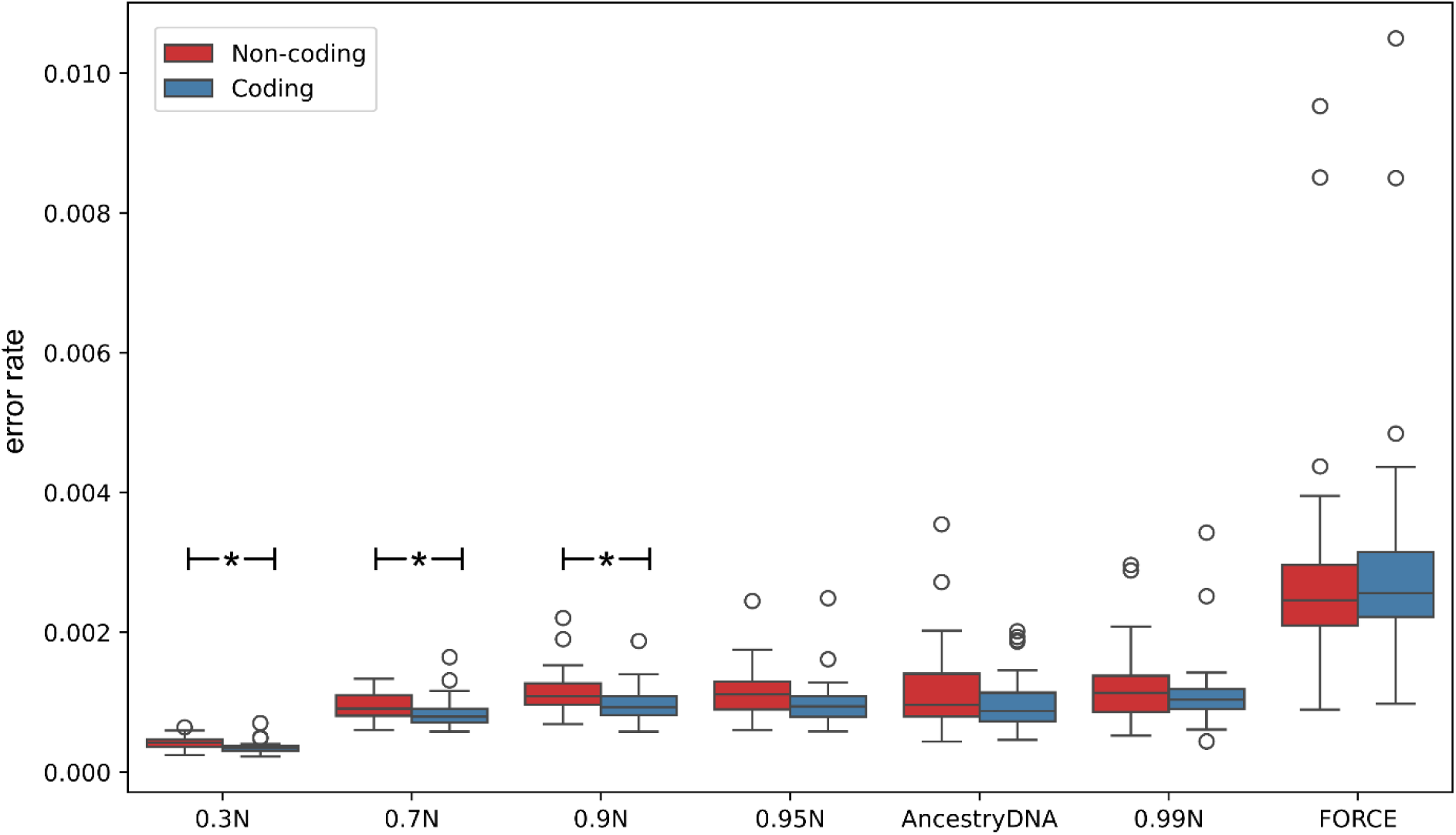
Imputation error rate distribution of SNP genotypes in non-coding (red) and coding (blue) regions of chromosome 16 over all 31 samples for each preimputation dataset. Error rates were obtained after normalising to equal numbers of coding and non-coding region SNPs per sample. Statistical significance was observed for the 3 largest preimputation datasets 0.3N, 0.7N and 0.9N.

### 3.4 Size of preimputation dataset and abundance of phenotype in reference panel impact phenotype prediction accuracy

Finally, we aimed to analyse the effect of imputed datasets on the prediction accuracy of phenotypic traits and tested whether the prediction accuracy was correlated with the representation of the phenotype in the imputation reference panel. The HIrisPlex-S prediction model for predicting eye, skin and hair colour was used for this test. The phenotype predictions using the complete datasets of the 31 test and 2,473 reference panel samples were considered as the “truth” (Supplementary Figure 3), as a validation of the HIrisPlex-S prediction model was out of the scope of this study. As described previously, the 31 test samples were selected specifically because they represent the 31 different phenotypes among all 1000 Genomes Project phase 3 individuals and additionally represent different biogeographic clusters (Supplementary Figure 4). The 2,473 reference panel samples represent 25 out of the 31 different phenotypes.

After performing SNP genotype imputation in all 31 individuals using 7 preimputation datasets with different number of SNPs, we re-predicted the HIrisPlex-S phenotypes. Generally, the number of correctly predicted phenotypes decreased with decreasing number of SNPs in the preimputation dataset (Figure 4). As expected, the panels AncestryDNA and FORCE were exceptions to this observation, as they included 29 out of 39 and all 39 out of 39 of the HIrisPlex-S SNPs due to intentional SNP selection during panel generation.

**Figure 4:**
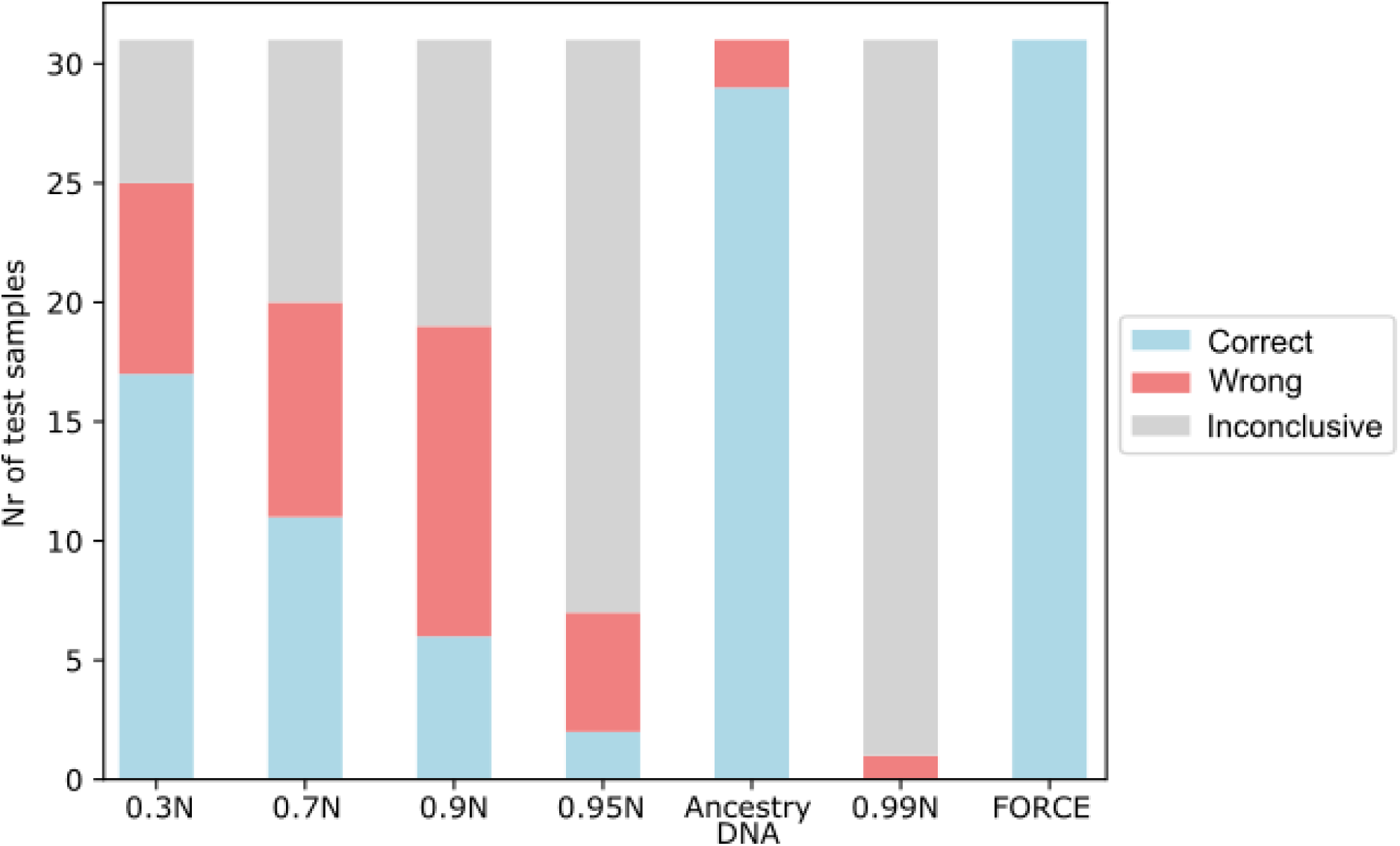
Number of correctly (blue) and incorrectly (red) predicted phenotypes (combination of eye, hair and skin colour) and inconclusive phenotypes (grey) of the 31 test samples in 7 preimputation datasets of decreasing number of SNPs prior to imputation. FORCE panel represents complete preimputation dataset without missing data.

Firstly, we investigated whether the biogeographic ancestry similarity between test and reference panel samples impacted the HIrisPlex-S prediction accuracy, since biogeographic ancestry strongly impacts imputation accuracy (1,7–10). Because shared population history increases the chance of shared LD blocks, similar populations or biogeographic ancestries are preferred for test and reference samples during imputation (26). For the current data no statistically significant correlation (Pearson correlation p-value=0.3003) was found between (A) the frequency of reference panel samples with the same biogeographic ancestry as the sample with a certain phenotype and (B) the sample’s phenotype prediction accuracy.

This concept was extended to test for the impact of frequency of shared phenotypes in the test and reference panel samples on the HIrisPlex-S prediction accuracy. The most frequently mispredicted phenotype across all preimputation datasets was the “true” phenotype of blue eyes/red hair/very pale skin, which was misidentified in all 6 imputation analyses (excluding FORCE). This phenotype was observed only 2 times in the reference panel. Interestingly, among the phenotypes that were most often correctly predicted was the very similar phenotype blue eyes/red hair/intermediate skin (13 observations in reference panel), as well as brown eyes/brown hair/dark to black skin (762 observations in reference panel). The latter was the most abundant phenotype in the reference panel (Supplementary Figure 3). A statistically significant correlation between abundance of a phenotype in the reference panel and how often this phenotype was correctly predicted in the imputation analyses was found (Pearson correlation= 0.456, p-value=9.932e-03).

However, these results should be considered with caution, as each of the 31 phenotypes were represented by only 1 sample and 7 different preimputation datasets of this sample. Further, the second most frequent phenotype (blue eyes/brown hair/dark to black skin) was represented in 388 individuals in the reference panel and was wrongfully predicted in 4 out of 6 imputation analyses. As this phenotype only differs in the eye colour from the most abundant phenotype described before, a prediction mix-up from blue to brown is not unexpected. Thus, very small allele alterations could result in different phenotypes, particularly for traits sensitive to misclassifications. The complex interplay between genetic markers to express phenotypic traits challenges the prediction model and thus erroneous imputations of few markers can have big implications in the predicted traits.

### 3.5 More lenient imputed SNP genotype calling thresholds cause most accurate trait predictions due to detrimental effect of missing SNP genotypes compared to incorrectly imputed SNP genotypes

To further explore the impact of the SNP genotype imputation on the prediction of individual traits, we split the HIrisPlex-S SNPs into those that encode for the 3 different traits: 6 eye colour-defining SNPs, 22 hair colour-defining SNPs (2 of these are missing in the 1000 Genomes Project dataset), and 36 skin colour-defining SNPs. For the following analysis, the FORCE panel was removed since it contains all HIrisPlex-S SNPs. We applied 6 genotype probability thresholds (0.99, 0.95, 0.9, 0.85, 0.5 and 0.2) on the imputed HIrisPlex-S SNP genotypes for the 6 different preimputation datasets. For the resulting imputed datasets, we then calculated the differences in allele dosage compared to the complete (or true) dataset. These allele dosage differences and information on whether the trait was predicted correctly, incorrectly or was not determined, were presented in Supplementary Figures 5 to 10.

We observed that for stricter (i.e., higher) genotype probability thresholds, overall fewer HIrisPlex-S SNPs were imputed with higher fractions of correctly imputed SNP genotypes (Figure 1, Supplementary Figures 5 to 10). This directly translates to the traits the SNPs predict, as the number of undetermined trait predictions decreases with more lenient (i.e., lower) genotype probability thresholds (Figure 5; grey line). Instead, more traits are correctly predicted (Figure 5; blue line).

**Figure 5:**
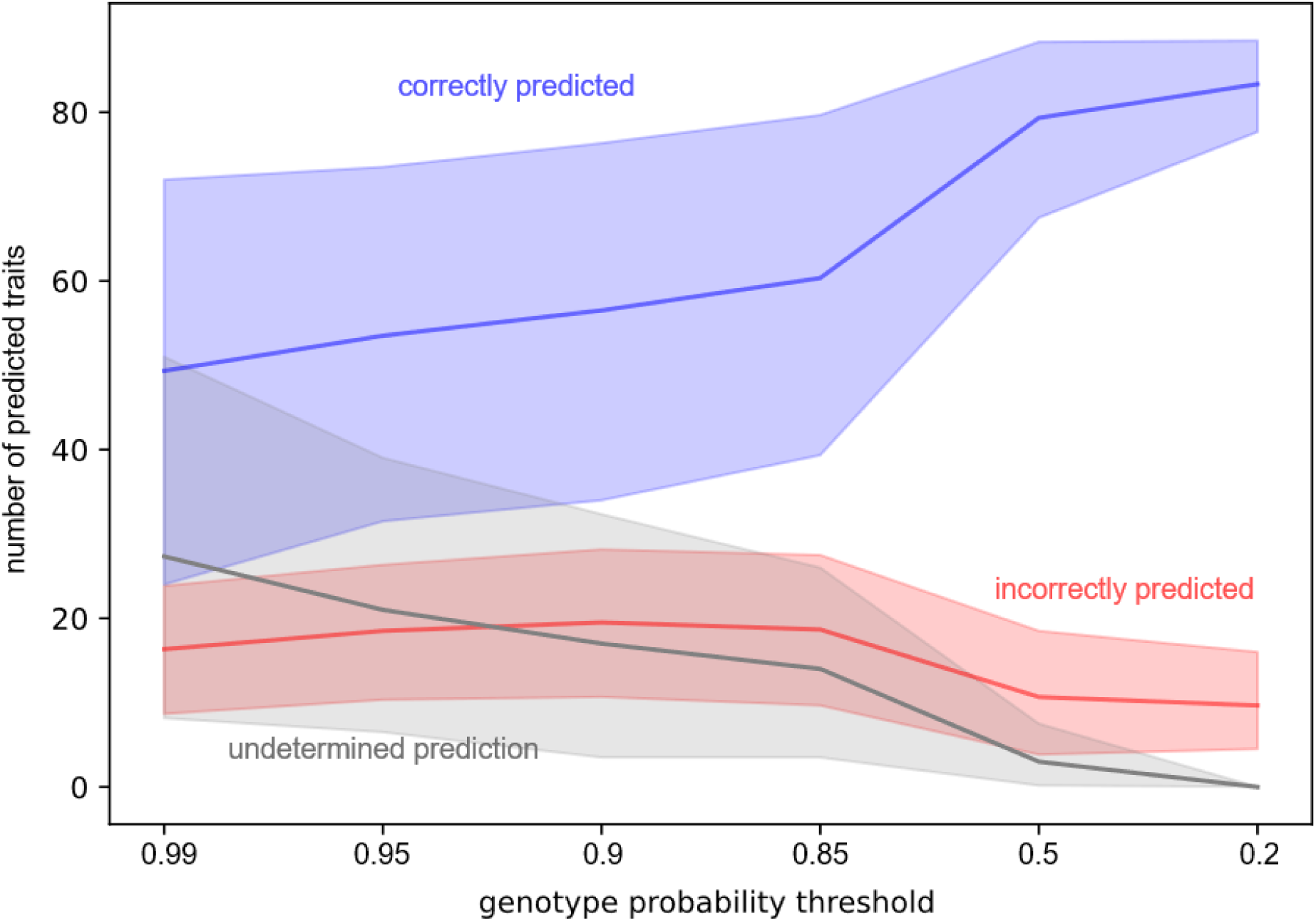
Number of correctly (blue), incorrectly (red) and undetermined (grey) trait predictions over all 31 samples. The lines represent the total number of traits of the 31 individuals averaged over the 6 different preimputation datasets (30% missing data to 99% missing data). The areas present 95% confidence intervals of the preimputation datasets. The trait prediction performance is presented for 6 different genotype probability thresholds with decreasing strictness from 0.99 to 0.2.

From strict to moderately strict probability thresholds (0.99≥t≥0.85), the incorrectly predicted traits increase as well (Figure 5; red line), most likely due to an unfavourable interaction of missing SNPs and incorrectly predicted SNPs. For t<0.85, the number of missing SNP genotypes decreases sufficiently, so that the information gained from available SNPs outweighs the negative implications from incorrectly imputed SNP genotypes. This particularly affects preimputation datasets with fewer initial SNPs compared to datasets with more initial SNPs (Supplementary Figure 11). Here, it becomes evident that a more lenient genotype probability threshold is most beneficial to improve the performance of the trait prediction.

Samples with correctly predicted traits tend to have lower rates of missing genotypes of the encoding SNPs compared to samples with incorrectly predicted traits (Supplementary Figure 12). However, the highest rates of missing genotypes were found when the trait could not be predicted. Multinomial logistic regressions were used to model the relationship of the 2 independent variables, missing genotypes and allele error rates, on the 3 prediction outcomes, i.e., correctly, incorrectly, and not determined trait predictions. For stricter imputation call thresholds t>0.5, missing SNP genotypes significantly increased misclassifications as “undetermined”, while also strongly impairing correct predictions. In comparison to missing genotypes, allele error rates had a weaker impact on the 3 possible prediction outcomes. For more lenient imputation call thresholds t≤0.5, the impact of missing genotypes on the prediction outcomes decreased down to no impact for t=0.2 with no missing data.

In general, the reason for incorrect trait predictions are usually not incorrectly imputed SNP genotypes, but rather the lack of represented SNP genotypes. Thus, we show that it is a misconception that stricter imputation criteria are necessary for reliable trait prediction. Instead, missing SNP genotypes have the most detrimental effect on trait prediction. However, this observation is specific to the HIrisPlex-S prediction model and might differ for other prediction tools.

## 4. Conclusion

In the current study, we investigated the performance of SNP genotype imputation with a focus on phenotype-correlated SNPs, which are commonly of interest upon SNP genotype imputation, but lack investigation in this context. We were able to corroborate previous findings for general SNPs on the impact of number and linkage of SNPs and MAFs on the imputation call and error rates.

Despite the higher error rates of phenotype-associated SNPs compared to randomly selected SNPs, a lenient genotype threshold enabled mostly correct trait predictions for the HIrisPlex-S prediction model. However, this might differ for other prediction models and thus highlights the importance of testing the impact of SNP genotype imputation criteria on the prediction model performance to gain the most reliable results.

We provide novel insights on the importance of genotyped or imputed SNP genotypes encoding a trait in the prediction model over erroneous SNP genotypes and prove the advantage of panels with highly linked SNPs for prediction models. These findings lay the groundwork for applying genotype imputation for forensic DNA phenotyping to provide investigative leads that can be combined with other scientific methods when repetitions of resource-intensive in-vitro experiments are not possible or successful.

