## Supplementary Figures for "Optimising Genotype Imputation for Precise Genetic Association in Forensic Phenotype Prediction and Trait Studies"

for

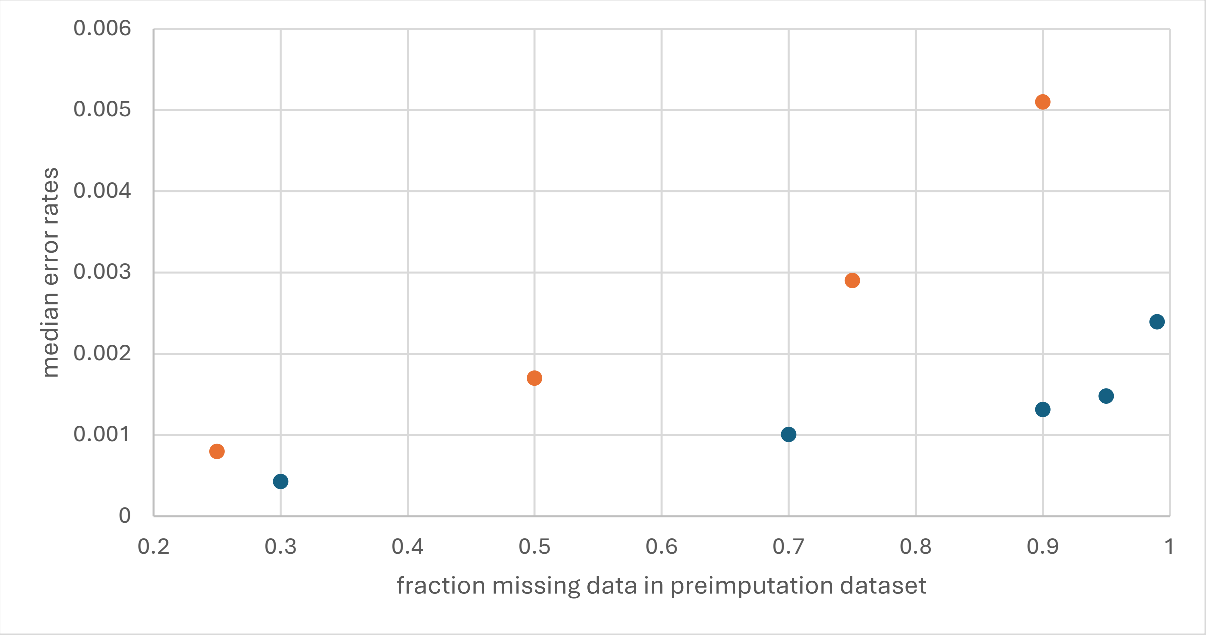


Supplementary Figure 1: Median error rates for SNP genotype imputations (≥0.95 genotype probability threshold) of increasing fractions of missing data in the AllSNPs preimputation datasets in the current study (blue) and Tillmar et al. 2024 (orange).

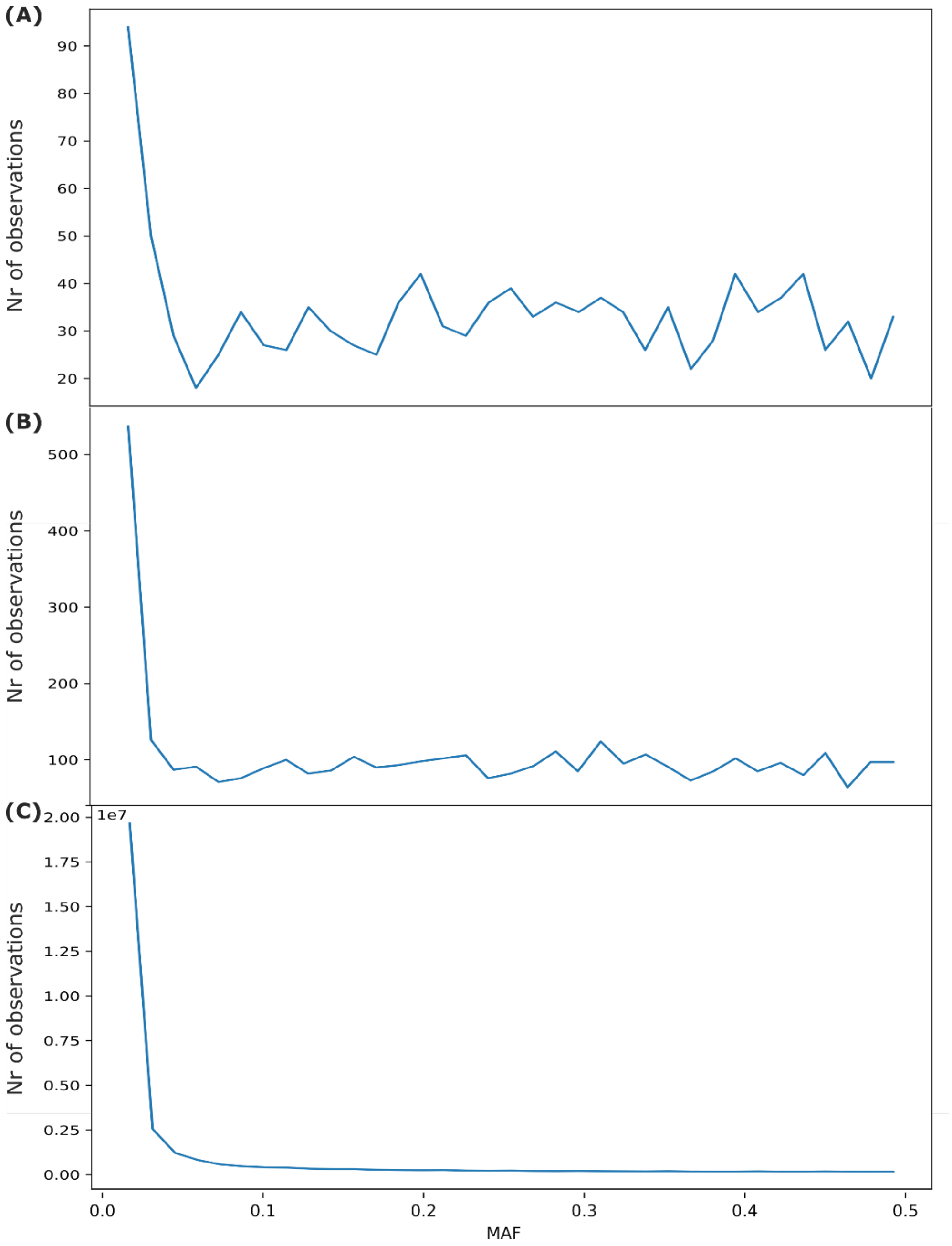


Supplementary Figure 2: Distribution of minor allele frequencies (MAF) for (A) PredSNPs, (B) PhenoSNPs and (C) AllSNPs.

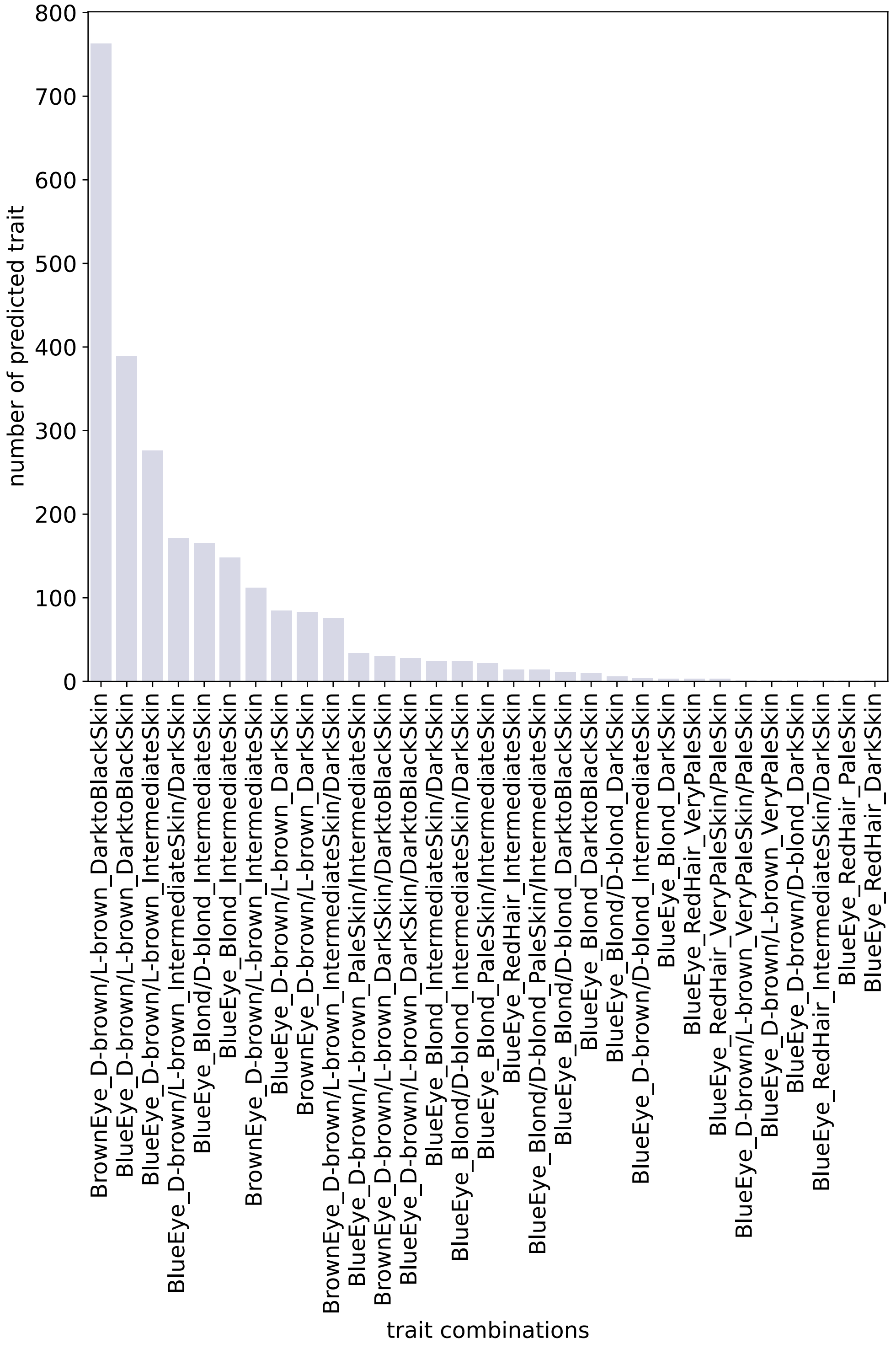


Supplementary Figure 3: Numbers of individuals from 1000 Genomes Project phase 3 data of each HIrisiplex-S-predicted phenotype using all 39 available SNPs.

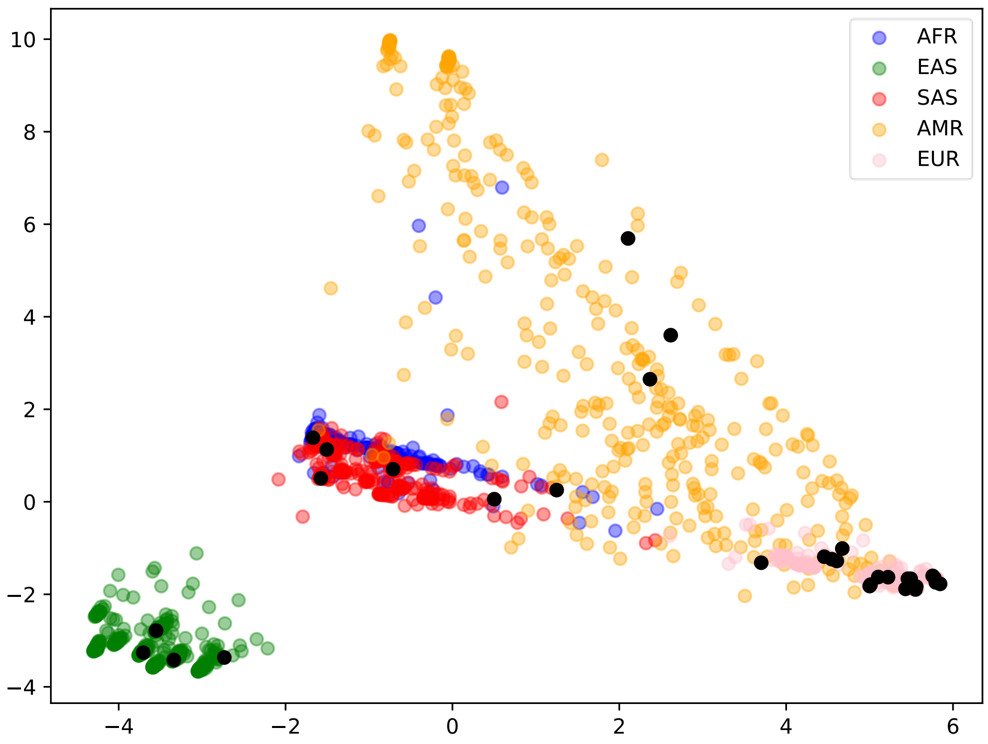


Supplementary Figure 4: Principal Component Analysis based on the three HIrisPlex-S prediction traits and STRUCTURE cluster coefficients of the 2,473 reference panel samples coloured according to metapopulation information from 1000 Genomes Project metadata and 31 test samples coloured in black.

**0.99 genotype probabilty threshold**

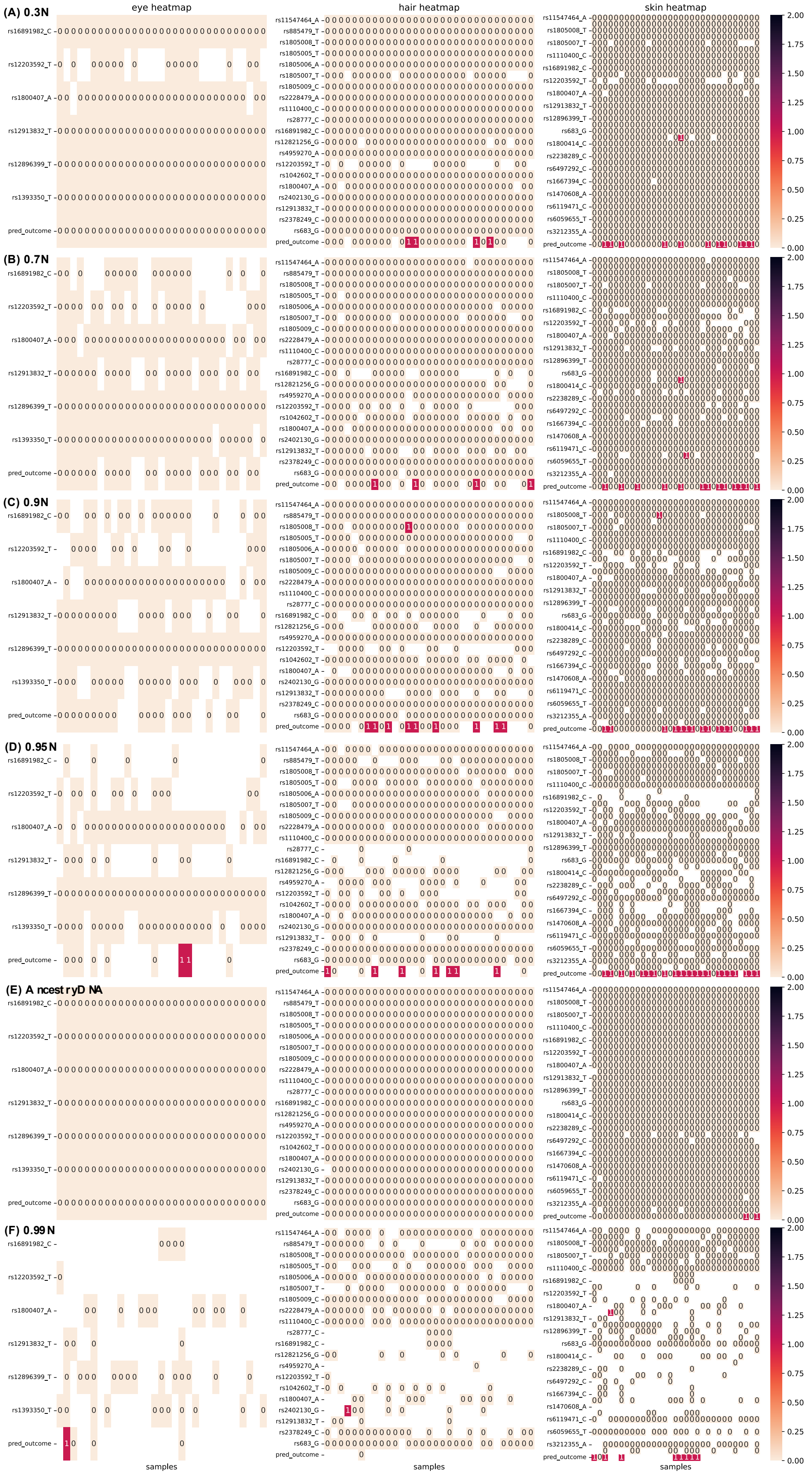

Supplementary Figure 5: Heatmaps of allele dosage differences (ranging from 0 to 2 or unknown) between complete dataset and imputed SNPs (0.99 genotype probability threshold) for preimputation datasets (A) 0.3N to (F) 0.99N. The analysed SNPs represented in rows are the 6 eye colour prediction SNPs (column 1), 22 hair colour prediction SNPs (column 2), and 36 skin colour prediction SNPs (column 3) implemented in the HIrisPlex-S prediction tool. The columns present the 31 investigated samples. The last row of each subplot shows whether the sample's trait was predicted correctly (0), wrongfully (1) or not at all (blank) when compared to the complete dataset.

#### 0.95 genotype probabilty threshold

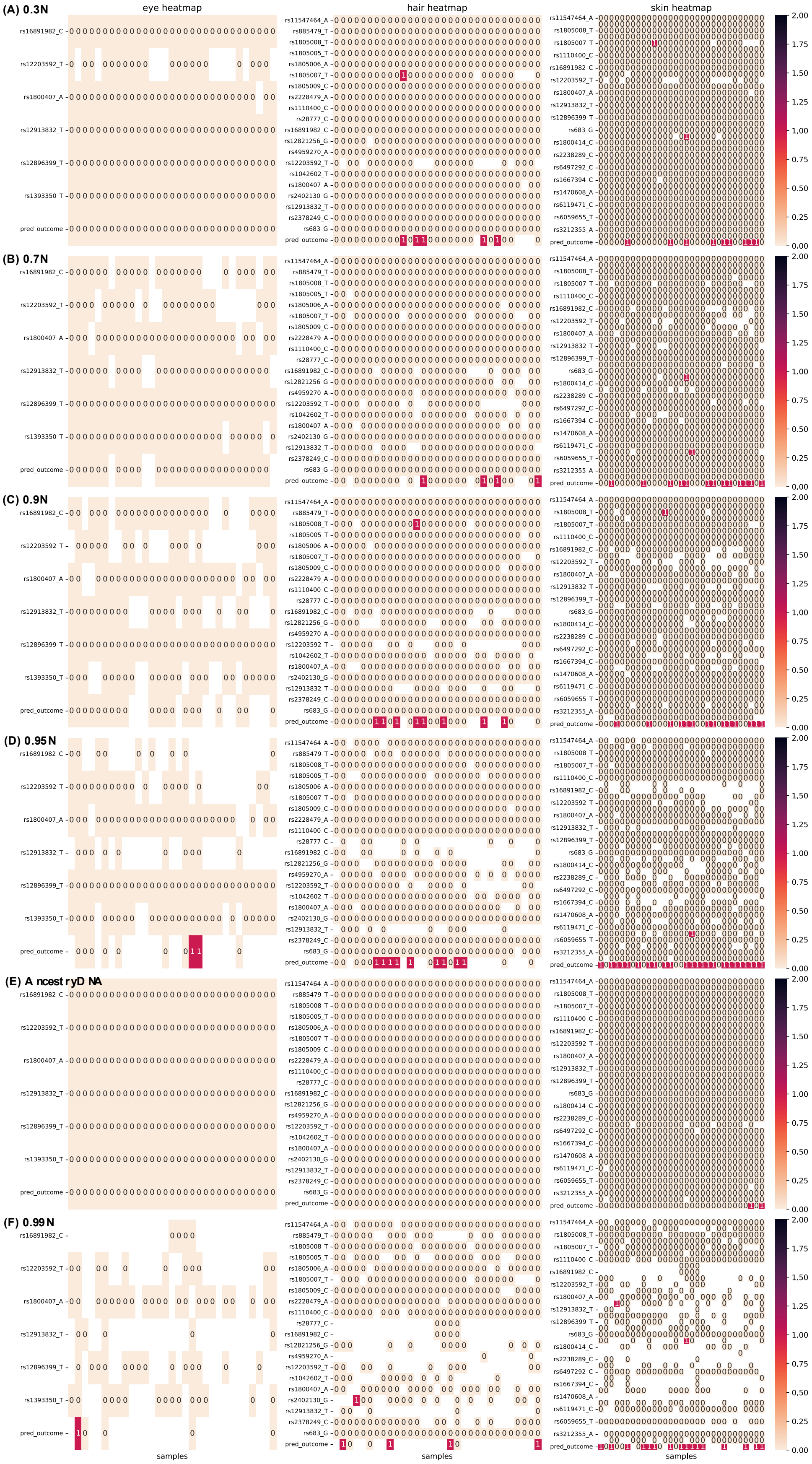

Supplementary Figure 6: Heatmaps of allele dosage differences (ranging from 0 to 2 or unknown) between complete dataset and imputed SNPs (0.95 genotype probability threshold) for preimputation datasets (A) 0.3N to (F) 0.99N. The analysed SNPs represented in rows are the 6 eye colour prediction SNPs (column 1), 22 hair colour prediction SNPs (column 2), and 36 skin colour prediction SNPs (column 3) implemented in the HIrisPlex-S prediction tool. The columns present the 31 investigated samples. The last row of each subplot shows whether the sample's trait was predicted correctly (0), wrongfully (1) or not at all (blank) when compared to the complete dataset.

#### 0.9 genotype probabilty threshold

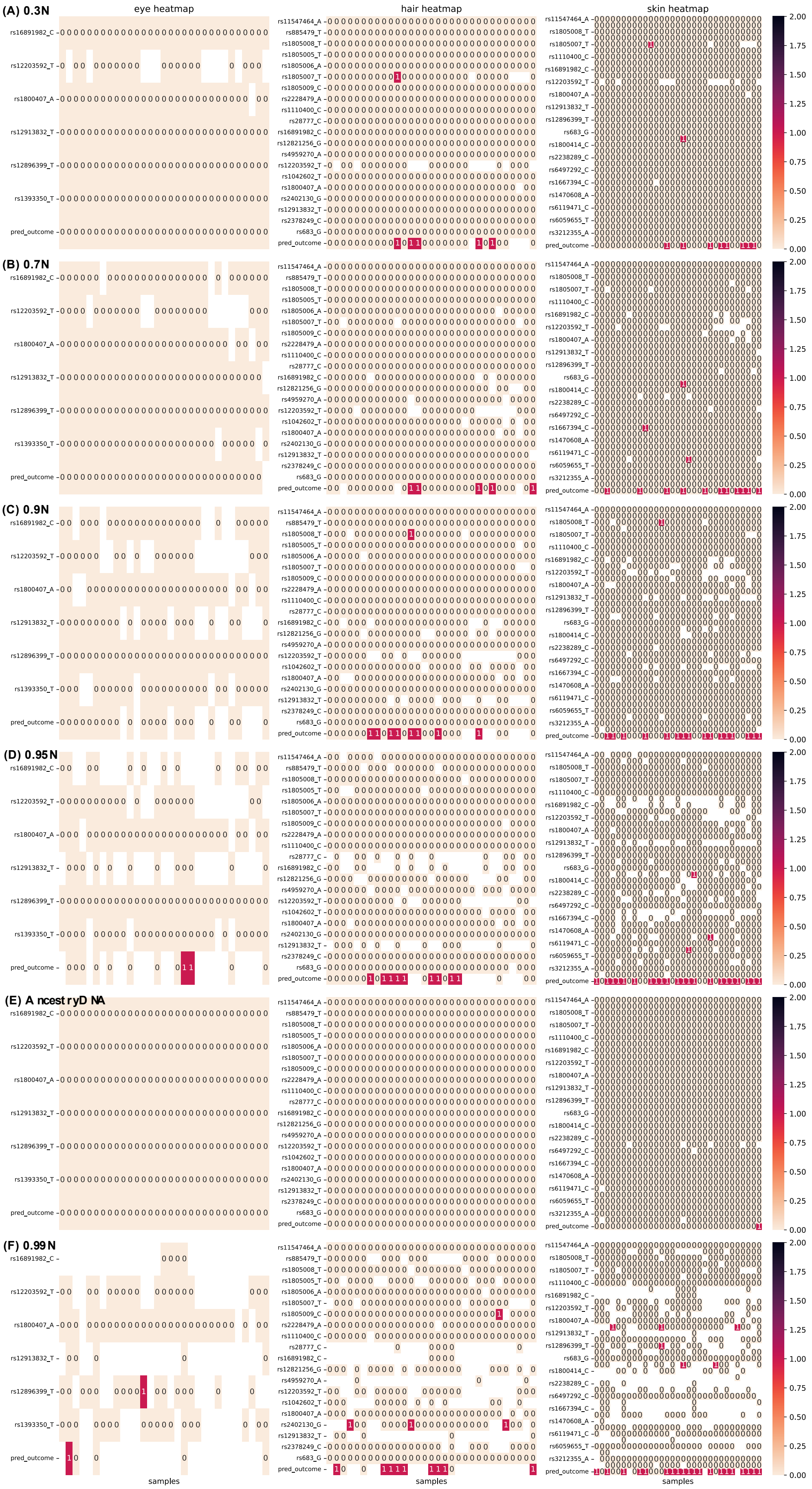

Supplementary Figure 7: Heatmaps of allele dosage differences (ranging from 0 to 2 or unknown) between complete dataset and imputed SNPs (0.9 genotype probability threshold) for preimputation datasets (A) 0.3N to (F) 0.99N. The analysed SNPs represented in rows are the 6 eye colour prediction SNPs (column 1), 22 hair colour prediction SNPs (column 2), and 36 skin colour prediction SNPs (column 3) implemented in the HIrisPlex-S prediction tool. The columns present the 31 investigated samples. The last row of each subplot shows whether the sample's trait was predicted correctly (0), wrongfully (1) or not at all (blank) when compared to the complete dataset.

#### 0.85 genotype probabilty threshold

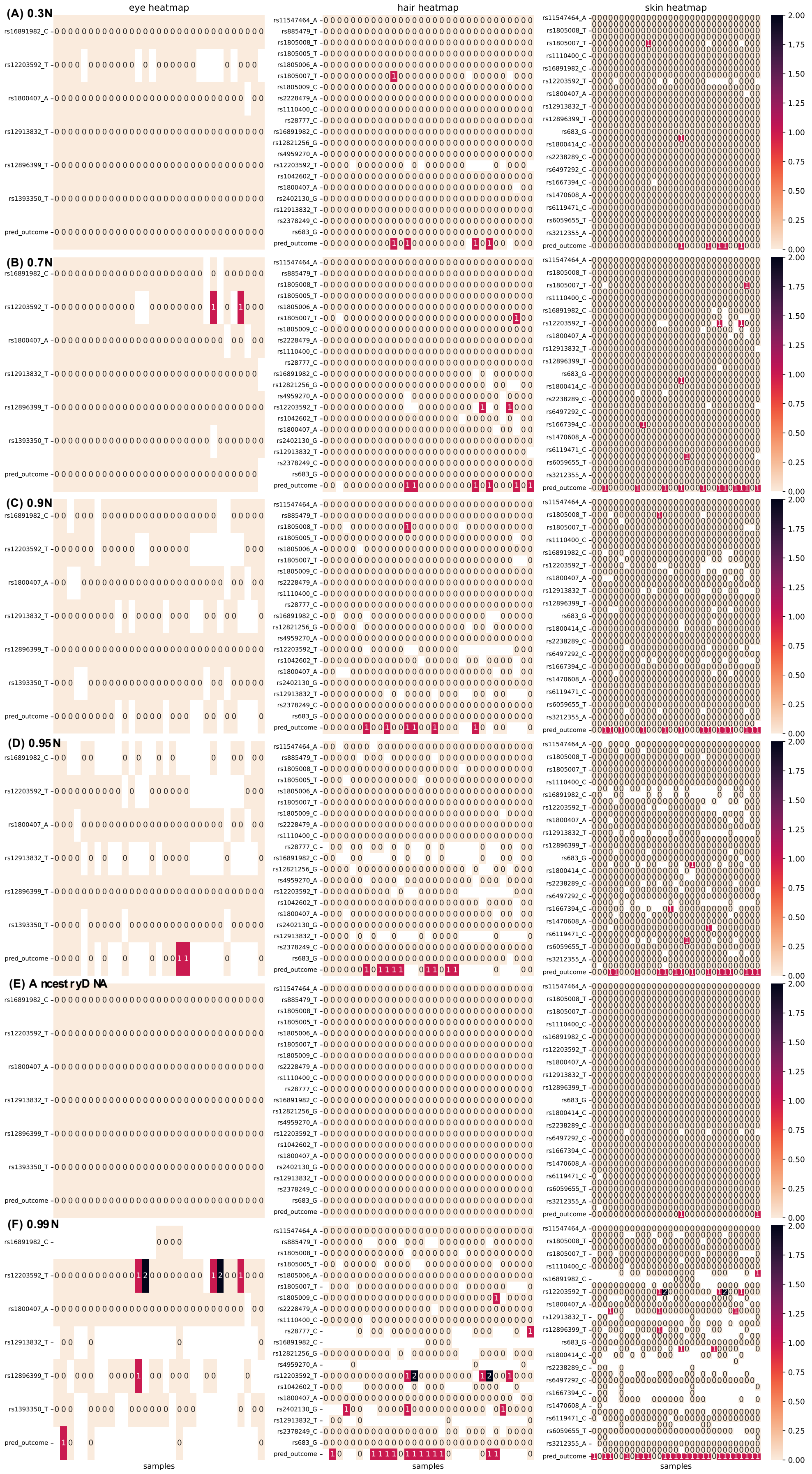

Supplementary Figure 8: Heatmaps of allele dosage differences (ranging from 0 to 2 or unknown) between complete dataset and imputed SNPs (0.85 genotype probability threshold) for preimputation datasets (A) 0.3N to (F) 0.99N. The analysed SNPs represented in rows are the 6 eye colour prediction SNPs (column 1), 22 hair colour prediction SNPs (column 2), and 36 skin colour prediction SNPs (column 3) implemented in the HIrisPlex-S prediction tool. The columns present the 31 investigated samples. The last row of each subplot shows whether the sample's trait was predicted correctly (0), wrongfully (1) or not at all (blank) when compared to the complete dataset.

**0.5 genotype probabilty threshold**

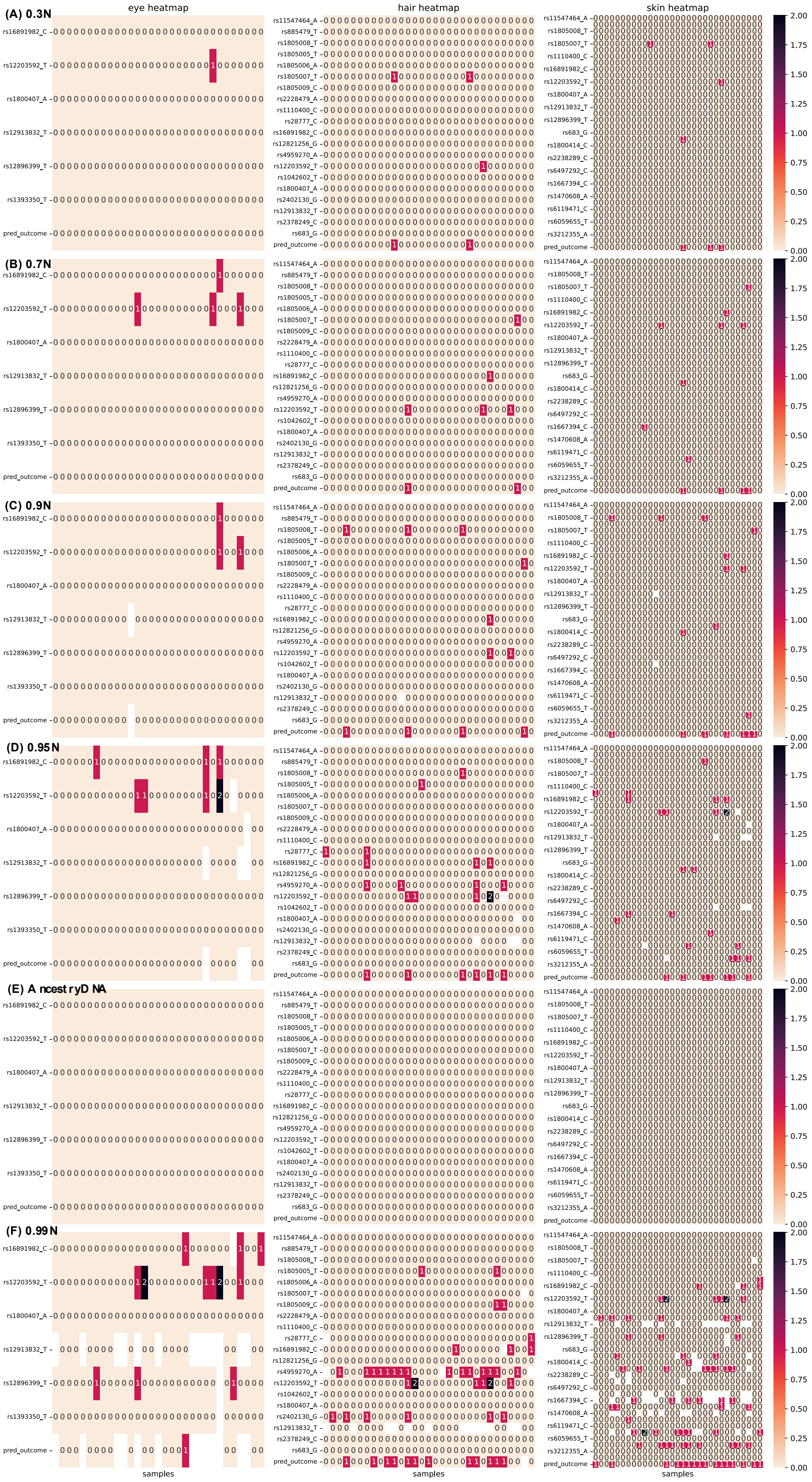

Supplementary Figure 9: Heatmaps of allele dosage differences (ranging from 0 to 2 or unknown) between complete dataset and imputed SNPs (0.5 genotype probability threshold) for preimputation datasets (A) 0.3N to (F) 0.99N. The analysed SNPs represented in rows are the 6 eye colour prediction SNPs (column 1), 22 hair colour prediction SNPs (column 2), and 36 skin colour prediction SNPs (column 3) implemented in the HIrisPlex-S prediction tool. The columns present the 31 investigated samples. The last row of each subplot shows whether the sample's trait was predicted correctly (0), wrongfully (1) or not at all (blank) when compared to the complete dataset.

### 0.2 genotype probabilty threshold

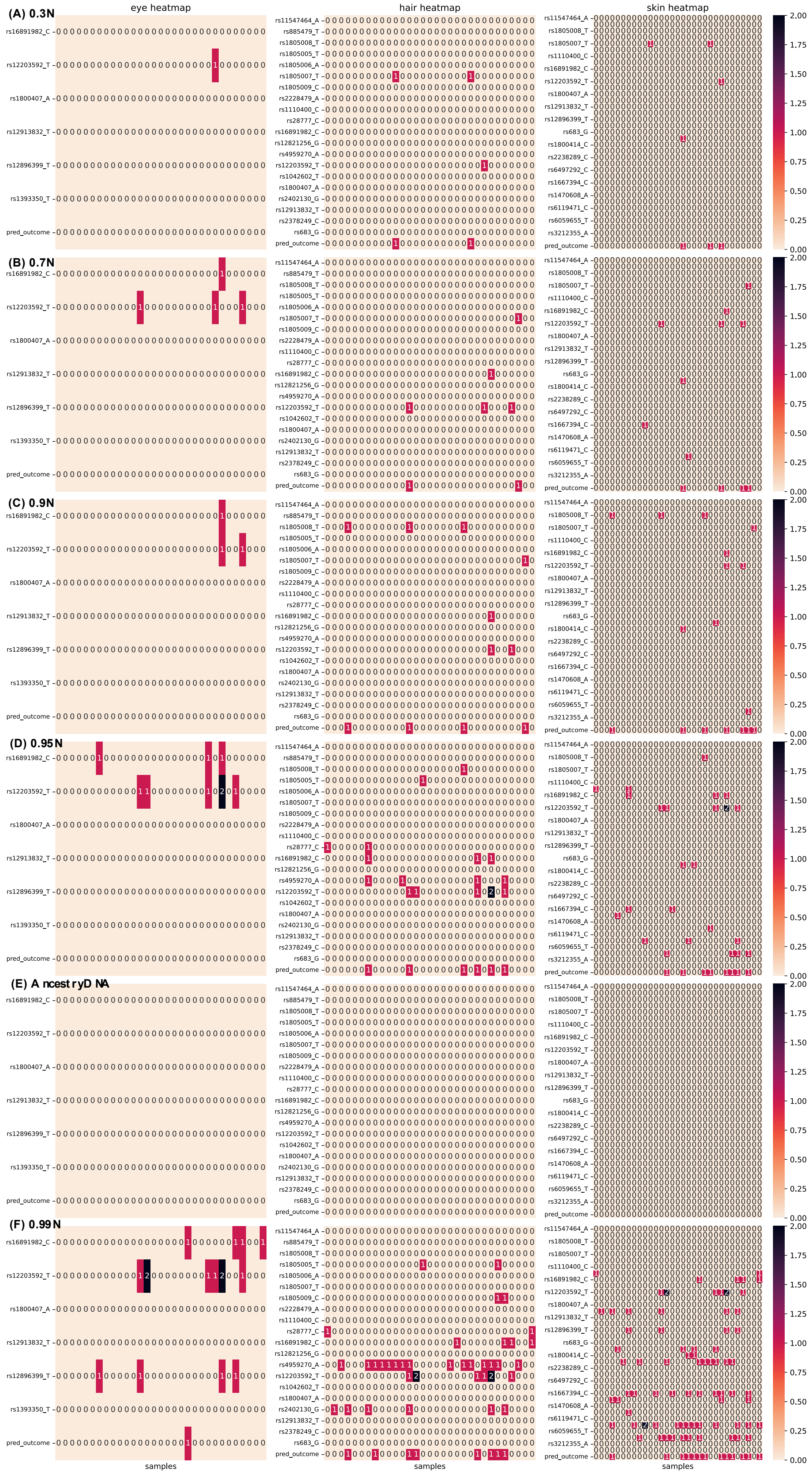

Supplementary Figure 10: Heatmaps of allele dosage differences (ranging from 0 to 2 or unknown) between complete dataset and imputed SNPs (0.2 genotype probability threshold) for preimputation datasets (A) 0.3N to (F) 0.99N. The analysed SNPs represented in rows are the 6 eye colour prediction SNPs (column 1), 22 hair colour prediction SNPs (column 2), and 36 skin colour prediction SNPs (column 3) implemented in the HIrisPlex-S prediction tool. The columns present the 31 investigated samples. The last row of each subplot shows whether the sample's trait was predicted correctly (0), wrongfully (1) or not at all (blank) when compared to the complete dataset.

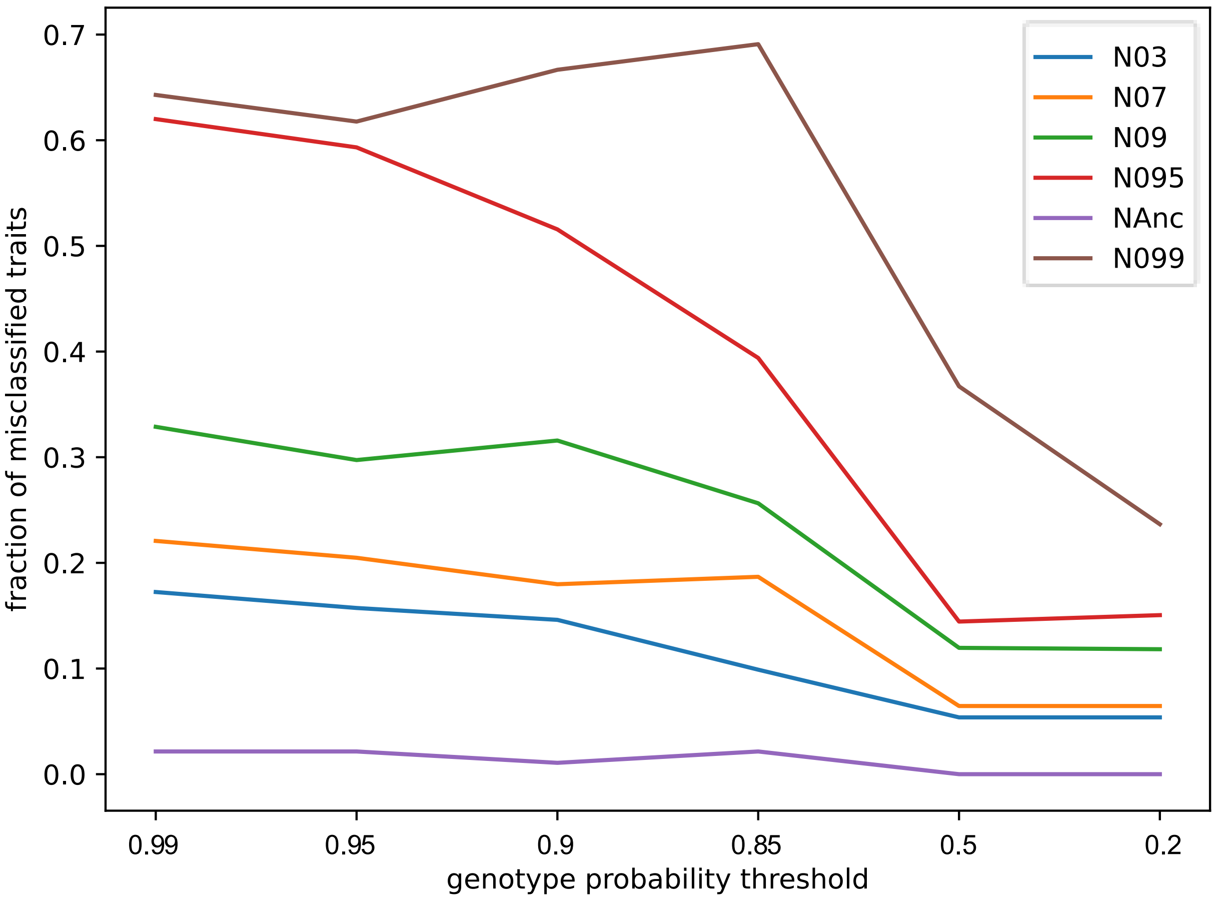


Supplementary Figure 11: Fraction of misclassified traits out of all predicted traits over all 31 individuals depending on six varying genotype probability thresholds ranging from 0.99 to 0.2 (x axis) for six difference preimputation datasets (fraction of missing data ranging from 0.3 to 0.99).

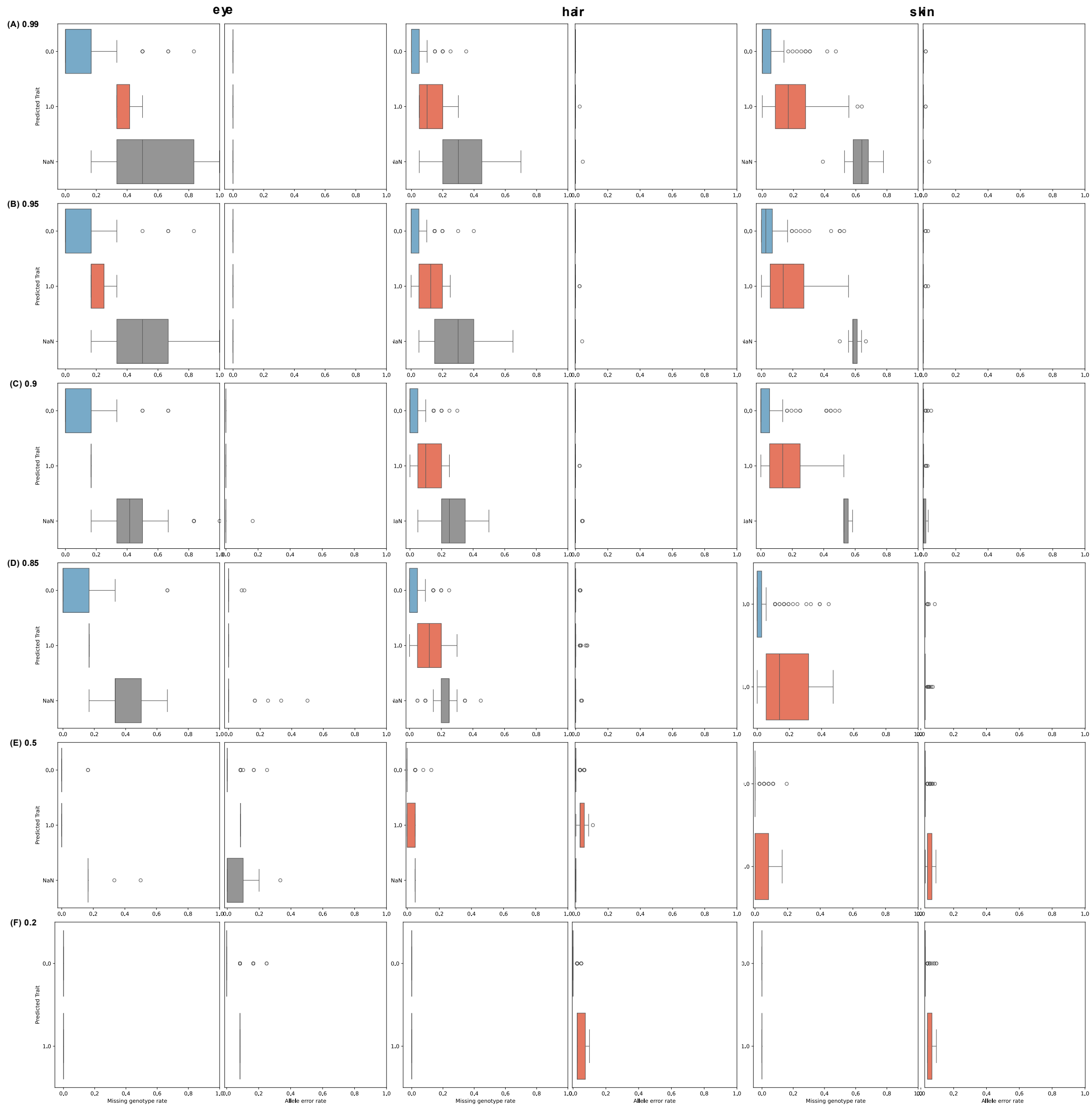


Supplementary Figure 12: Distribution of per-sample and post-imputation missing genotype rates (left x axes) and allele error rates (right x axes) for SNP genotypes incorporated in predicting eye colour (column 1), hair colour (column 2) and skin colour (column 3). The corresponding individual's trait was predicted correctly (prediction trait: 0; blue), incorrectly (prediction trait: 1; red) or was undetermined (prediction trait: NaN; grey). Each row represents imputations with different genotype probability thresholds from (A) 0.99 to (F) 0.2.
